# Prediction of antibiotic resistance mechanisms using a protein language model

**DOI:** 10.1101/2024.05.04.592288

**Authors:** Kanami Yagimoto, Shion Hosoda, Miwa Sato, Michiaki Hamada

## Abstract

**Motivation:** Antibiotic resistance has emerged as a major global health threat, with an increasing number of bacterial infections becoming difficult to treat. Predicting the underlying resistance mechanisms of antibiotic resistance genes (ARGs) is crucial for understanding and combating this problem. However, existing methods struggle to accurately predict resistance mechanisms for ARGs with low similarity to known sequences and lack sufficient interpretability of the prediction models.

**Results:** In this study, we present a novel approach for predicting ARG resistance mechanisms using Protein-BERT, a protein language model based on deep learning. Our method outperforms state-of-the-art techniques on diverse ARG datasets, including those with low homology to the training data, highlighting its potential for predicting the resistance mechanisms of unknown ARGs. Attention analysis of the model reveals that it considers biologically relevant features, such as conserved amino acid residues and antibiotic target binding sites, when making predictions. These findings provide valuable insights into the molecular basis of antibiotic resistance and demonstrate the interpretability of protein language models, offering a new perspective on their application in bioinformatics.

**Availability:** The source code is available for free at https://github.com/hmdlab/ARG-BERT. The output results of the model are published at https://waseda.box.com/v/ARG-BERT-suppl.

## 1 Introduction

The prevalence of antibiotic-resistant bacteria is increasing and poses a significant threat to global public health. If no action is taken, the annual number of deaths resulting from such infections is expected to reach 10 million by 2050 (O’Neill, 2016). Thus, it is imperative to urgently address the spread of antibiotic resistance. Recent global and national action plans prioritize the One Health approach (Organization *et al*., 2015), recognizing the relationships of human, animal, and environmental health. Resistant bacteria are found in various niches across these domains, creating a reservoir of antibiotic-resistance genes (ARGs) (Kim and Cha, 2021). These reservoirs can potentially amplify the harm caused by antibiotic resistance through horizontal gene transfer (Cox and Wright, 2013). Investigating ARG reservoirs is crucial in mitigating the damage caused by antibiotic resistance, which requires strategies for identifying ARGs and predicting and analyzing resistance mechanisms (Alcock *et al*., 2023).

Current approaches for predicting resistance mechanisms can be categorized into two main types. The first and most common approach, exemplified by CARD-RGI (Alcock *et al*., 2023; Hendriksen *et al*., 2019), relies on sequence homology search tools. These tools are employed to identify sequences similar to the input sequences from the ARG reference database. The advantage of these methods is a lower false-positive rate due to the similarity score cut-off. However, it is crucial to recognize that the cut-off similarity score restricts the applicability of these methods to ARGs with low similarity to sequences in the reference database, potentially leading to an increased false-negative rate (Arango-Argoty *et al*., 2018; Hendriksen *et al*., 2019; Li *et al*., 2021). The second approach involves the application of deep learning models. For instance, HMD-ARG (Li *et al*., 2021) has employed a convolutional neural network (CNN), while LM-ARG (Ahmed *et al*., 2022) has used embedding from pLM(protein language models). Although these models can accurately predict sequences with low similarity to reference databases, they do not provide any interpretation of the predicted resistance mechanisms (Li *et al*., 2021). In summary, there is currently no resistance mechanism prediction method that achieves both scalability to sequences dissimilar to the reference database and interpretability of the results. Therefore, predictive models of resistance mechanisms incorporating these features are essential to tackle the challenge of antibiotic resistance. Bidirectional Encoder Representations from Transformers (BERT) (Devlin *et al*., 2019), a deep learning model in natural language processing, may offer accuracy and interpretability. BERT-based pLMs, pre-trained on large protein datasets, have demonstrated improved performance and interpretability compared to existing deep learning models (Elnaggar *et al*., 2021; Brandes *et al*., 2022; Vig *et al*., 2020). The interpretability is attributed to attention, which identifies biological features contributing to the task (Zhou *et al*., 2023; Vig *et al*., 2020; Lee *et al*., 2023; Yamada and Hamada, 2022; Brandes *et al*., 2022).

In this study, we propose a novel method for predicting resistance mechanisms of ARGs using ProteinBERT (Brandes *et al*., 2022), a pLM pre-trained on a large protein dataset (See Figure 1 for an overview of this study.). Our model performed as well as, or better than, existing models for ARG datasets containing sequences with low similarity to reference sequences. Through attention analysis, we demonstrated the interpretability of our model by identifying a correlation between attention and amino acid conservation and a concentration of attention on target binding sites and conserved regions involved in antibiotic resistance. Our proposed approach presents a promising solution to the challenges associated with antibiotic resistance.

**Fig. 1:**
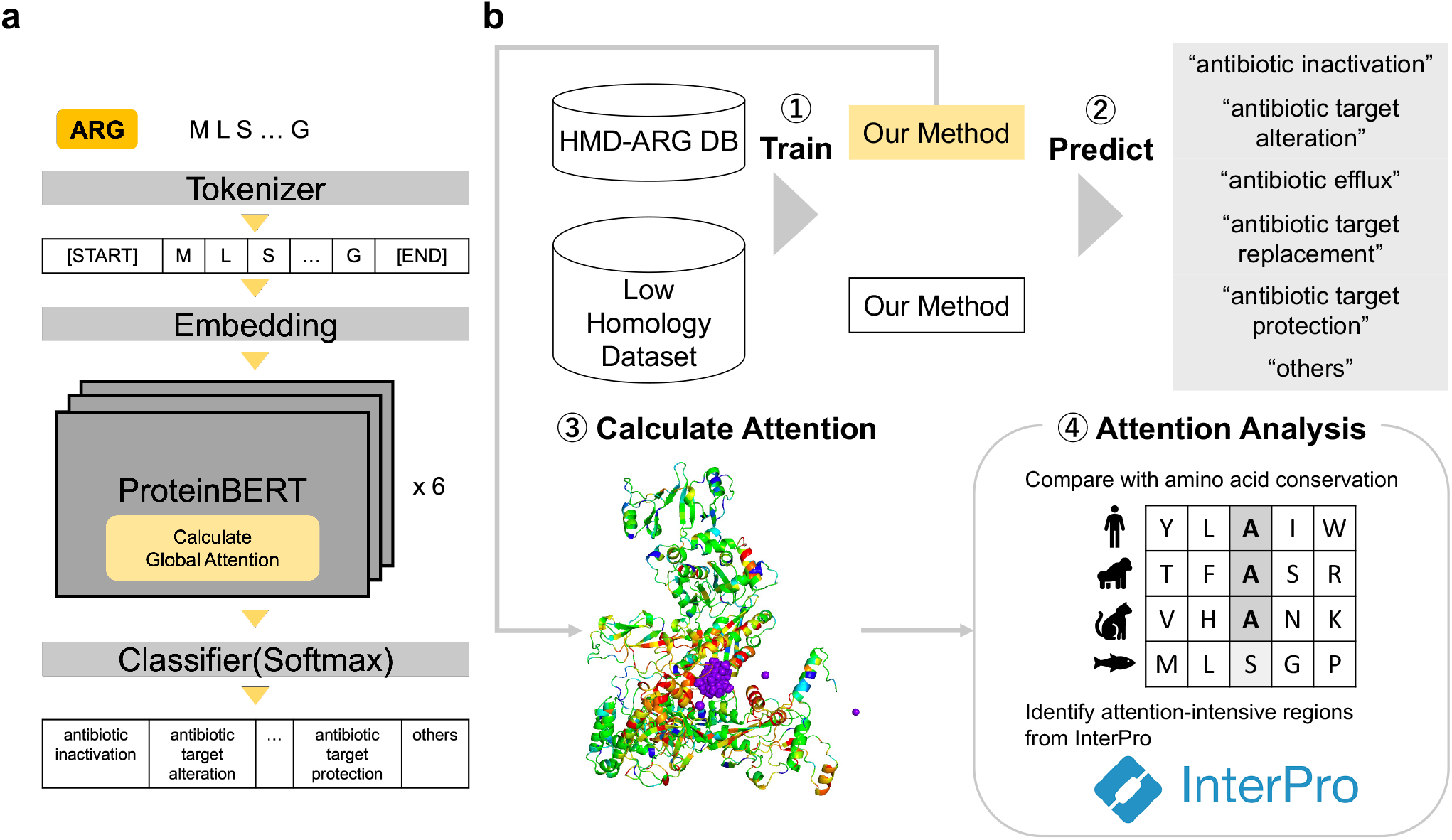
Overview of this study. (a) The architecture of the proposed model for predicting antibiotic resistance gene (ARG) resistance mechanisms. The model, based on ProteinBERT, a BERT-based protein language model (pLM), includes an input layer for ARG sequences, and an output layer for predicting six resistance mechanism labels: antibiotic efflux, antibiotic inactivation, antibiotic target alteration, antibiotic target protection, antibiotic target replacement, and others. (b) The model was fine-tuned using the HMD-ARG dataset and a custom-built low homology dataset. Subsequently, attention analysis was performed on the fine-tuned model (with HMD-ARG dataset) to interpret its predictions, suggesting that the proposed method takes into account biologically relevant features, such as antibiotic-binding sites and evolutionarily conserved regions, when making predictions.

## 2 Materials and Methods

### 2.1 Dataset collection

In this study, two datasets were utilized to train and evaluate the model as follows.

#### 2.1.1 HMD-ARG dataset

The first dataset, the HMD-ARG dataset, was created by Li *et al*. (2021) and is currently the largest available database, integrating seven publicly available ARG databases. The DNA sequences were converted to protein sequences, and duplicate sequences were removed before merging the seven ARG databases, resulting in a total of 24,082 ARGs. Each ARG was then annotated with labels corresponding to antibiotic efflux, antibiotic inactivation, antibiotic target alteration, antibiotic target protection, antibiotic target replacement, and other resistance mechanisms. The annotation of resistance mechanism labels was based on the CARD (Alcock *et al*., 2023) ontology, using BLASTP (Altschul *et al*., 1990) or literature review. The complete dataset, along with the number of sequences corresponding to each resistance mechanism, is presented in Supplementary Figure S1. See also the previous paper (Li *et al*., 2021) for details of this dataset.

To assess the performance using the datasets, we employed *stratified* 5-fold cross-validation (CV). Each dataset was divided into five separate groups, ensuring that the proportion of resistance mechanisms in each group was equal.

#### 2.1.2 Low homology dataset

To evaluate the performance of the proposed method on sequences with low similarity to known ARGs, we created a second dataset from the HMD-ARG dataset (Section 2.1.1), named the ‘Low Homology Dataset,’ which features reduced homology between training and test data. First, we employed CD-HIT (L *et al*., 2012) to cluster sequences with the same resistance mechanism and sequence identity greater than 0.4. Each cluster is regarded as an identical group. We divided the unclustered dataset into five groups and added sequences assigned to 5-folds. In our evaluation, we used the Group K-fold package in Scikit-learn (Pedregosa *et al*., 2011) for the clusters and mechanisms. This approach resulted in a maximum sequence identity of 0.4 between training and test data for sequences with the same resistance mechanism. The same procedure was then applied to create datasets with sequence identity thresholds of 0.5, 0.6, 0.7, 0.8, and 0.9.

### 2.2 A Pre-trained Protein Language Model

In this study, we employed ProteinBERT, a type of protein language model (pLM) pre-trained on protein sequences and Gene Ontology (GO) terms (Ashburner *et al*., 2000; Gene Ontology *et al*., 2023) that define the biological functions of genes. ProteinBERT consists of two components: one that processes GO terms as a *global* representation, 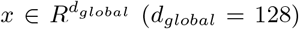, and another that processes amino acid sequence data as a *local* representation 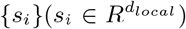, where *i* shows the position in a protein sequence. The global and local representations are connected via global attention (Section 2.6). In the pre-training step, the architecture was trained on 106 million protein sequences from UniProtKB/UniRef90 (Suzek *et al*., 2007) and 9, 000 GO terms by applying the same Masked Language Modeling (MLM) method as BERT. See Brandes *et al*. (2022) for the detailed model of ProteinBERT.

ProteinBERT offers several advantages as a pLM. First, it has fewer parameters than other pLMs and does not require training on large datasets, facilitating pre-training on non-redundant datasets and avoiding biased pre-training on specific protein families. Consequently, fine-tuning requires less time and memory. Second, pre-training can utilize information from the Gene Ontology to learn functional information about proteins.

### 2.3 Fine-tuning

We fine-tuned ProteinBERT to predict the resistance mechanism of each ARG (cf. Figure 1). To predict one of the six resistance mechanism labels, we first added a fully connected layer to the global representation output of ProteinBERT (described in the previous section). We then trained and tested the model using stratified (or stratified group) 5-fold cross-validation. Specifically, using the dataset divided in the way described in Section 2.1, we trained one group and tested the model on the remaining one. This process was repeated five times and the average results were reported. For improvement of performance, we followed the training approach proposed by Brandes *et al*. (2022). Training and evaluation were conducted using two NVIDIA A100 (80GB) GPUs, which took up to 12 minutes. See Supplementary Section S2.2 for the detailed fine-tuning procedure.

### 2.4 Comparison with baseline models

The performance of the proposed method was compared with that of LM-ARG, HMD-ARG, BLAST, and CARD-RGI. LM-ARG predicts the resistance mechanism by passing the embedding provided by ProtAlbert (Elnaggar *et al*., 2021), one of the pLMs, through the pooling and feed-forward layers (Ahmed *et al*., 2022). HMD-ARG is a resistance mechanism prediction model using 6-layer CNN, developed by Li *et al*. (2021). Although they predicted only sequences predicted as ARG at Level 0, we predicted all input sequences of resistance mechanisms. BLAST(Altschul *et al*., 1990) was used to predict by retrieving similar sequences from the test set, using the training data as a reference. The resistance mechanism was estimated using the hit sequence with the lowest e-value. CARD-RGI is a BLAST-based model that predicts resistance mechanisms based on bit scores, using The Comprehensive Antibiotic Resistance Database (CARD) (Alcock *et al*., 2020, 2023) as a reference. In the prediction made by BLAST and CARD-RGI, sequences with no similar sequence hit in reference sequences were classified as “others”.

### 2.5 Evaluation metrics

We evaluated the performance of the proposed method by employing indicators of Accuracy, Precision, Recall, and F1-score. For Precision, Recall, and F1-score, we considered the prediction of each resistance mechanism as a binary classification problem, distinguishing between the focused mechanism and all other mechanisms. Based on this, we calculated the True Positive (TP), True Negative (TN), False Positive (FP), and False Negative (FN) values for each resistance mechanism. These metrics were then computed using the following equations:

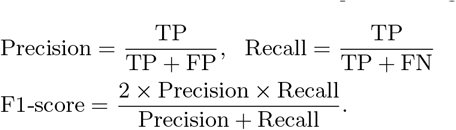

For Precision, Recall, and F1-score, we calculated the metrics for each resistance mechanism and then computed the macro average. Accuracy was calculated as the proportion of sequences for which the resistance mechanism was correctly predicted among all sequences in the dataset. We repeated this process for each iteration of 5-fold cross-validation and then obtained the average of all iterations.

### 2.6 Attention analysis

Attention value is computed in ProteinBERT to incorporate local representations into global representations. The value is a per-amino acid value that is calculated as *z*_1_, …, *z*_*L*_ in the following formula:

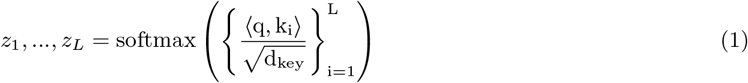

where 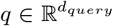 and 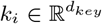 are the query and key vectors, respectively. The *q* and *k*_*i*_ are used for incorporating the global and local representation into global attention. They are calculated as *q* = tanh(*W*_*q*_*x*), *k*_*i*_ = tanh(*W*_*k*_*s*_*i*_) where *x* is the global representation, *s*_*i*_ is the local representation, and 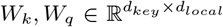 are the weight parameters.

To elucidate the biological significance of attention in the proposed method, we performed two analyses: (i) comparison with amino acid conservation (Section 2.6.1) and (ii) identification of functionally relevant domains where attention is concentrated (Section 2.6.2).

#### 2.6.1 Comparison with amino acid conservation

We examined the relationship between attention and a conservation score, which indicates the degree of conservation for each amino acid residue among the evolutionary-related bacteria. See Supplementary Section S2.3 for the details of computing the conservation score. First, we computed the Spearman correlation between attention and conservation scores. Second, we divided the residues into three groups according to their attention values, using the 33rd and 66th percentiles as thresholds. Finally, we assessed differences in the distribution of conservation scores between these groups using the Mann-Whitney U test (one-sided, greater) with Bonferroni correction (significance level p = 0.05/N, where N is the number of tests, N = 3).

#### 2.6.2 Identification of ‘attention-intensive regions’ and their biological function

We identified sequence regions where attention is concentrated and named them ‘attention-intensive regions’. We used InterProScan (Jones *et al*., 2014; Paysan-Lafosse *et al*., 2023) to identify specific sequence regions corresponding to families, domains, and functional sites conserved in the analyzed sequences. We then compared the distribution of attention in these regions with the whole sequence using the Mann-Whitney U test (one-sided, greater), and the significant regions are defined as ‘attention-intensive regions’.

We attempted to clarify the biological function of attention-intensive regions. Identified sequence regions were annotated with Gene Ontology terms by InterProScan(Jones *et al*., 2014). We performed GO enrichment analysis to infer the function of the attention-intensive sequence regions based on Gene Ontology (Gene Ontology *et al*., 2023). We used Fisher’s exact test (one-sided, greater) to determine whether each GO term was significantly enriched in attention-intensive regions. We restricted the sequence regions used in our analysis to those annotated with at least one GO term.

In the above, to determine significance, we applied the Bonferroni correction in all analyses (significance level p = 0.05/N, where N is the number of tests).

## 3 Results

### 3.1 Performance comparison

First, the performance of the proposed method was compared to that of LM-ARG, HMD-ARG, BLASTP, and CARD-RGI (cf. Section 2.4) using the HMD-ARG DB. The performance indicators of the proposed method, namely Accuracy, Precision, Recall, and F1-score, achieved 0.999, 0.943, 0.934, and 0.937 (Table 1, top). On the other hand, the existing methods with the best performance, LM-ARG and BLAST, were 0.980, 0.929, 0.902, 0.911, and 0.987, 0.910, 0.933, 0.919 respectively. Additionally, the detailed performances for each resistance mechanism are also shown in Supplementary Figure S3. (The ‘other’ resistance mechanism was excluded due to the small number of sequences.) All the methods achieved better performances for the resistance mechanisms with a large number of sequences than for those with a small number of sequences. In summary, when trained on the HMD-ARG DB, the proposed model performed slightly better than the best-performing existing methods, BLAST and LM-ARG.

**Table 1:** The comparison of performances for two datasets, HMD-ARG dataset and Low Homology Dataset. The bold values indicate the highest value for a combination of metric and dataset.

|  | Dataset | Accuracy | Precision | Recall | F1 Score |
| --- | --- | --- | --- | --- | --- |
| Proposed | HMD-ARG<br>DB | **0.999** | **0.943** | **0.934** | **0.937** |
| LM-ARG |  | 0.980 | 0.929 | 0.902 | 0.911 |
| HMD-ARG |  | 0.968 | 0.920 | 0.871 | 0.889 |
| BLAST |  | 0.987 | 0.910 | 0.933 | 0.919 |
| CARD-RGI |  | 0.910 | 0.803 | 0.885 | 0.784 |
| Proposed | Low | **0.927** | **0.869** | **0.871** | **0.866** |
| LM-ARG | Homology | 0.926 | 0.835 | 0.797 | 0.800 |
| BLAST | Dataset | 0.917 | 0.749 | 0.827 | 0.765 |

Next, we performed a performance comparison using the low homology dataset (Table1 bottom and Figure 2). The proposed method achieved an Accuracy of 0.927, Precision of 0.869, Recall of 0.871, and F1 score of 0.866, outperforming the other methods, which achieved values of 0.926, 0.835, 0.797, 0.800 (LM-ARG), and 0.917, 0.749, 0.827, 0.765 (BLAST). Furthermore, the proposed method was compared with LM-ARG and BLAST, which had performed well on the HMDARG-DB, on datasets with sequence identity thresholds ranging from 0.4 to 0.9. The proposed method notably surpassed LM-ARG and BLAST on the datasets with sequence identity thresholds of 0.4 and 0.5 (Figure 2). The performances for each mechanism are shown in Supplementary Figure S4. These results suggest that the proposed model accurately predicts resistance even for ARGs with low sequence similarity to known ARGs.

**Fig. 2:**
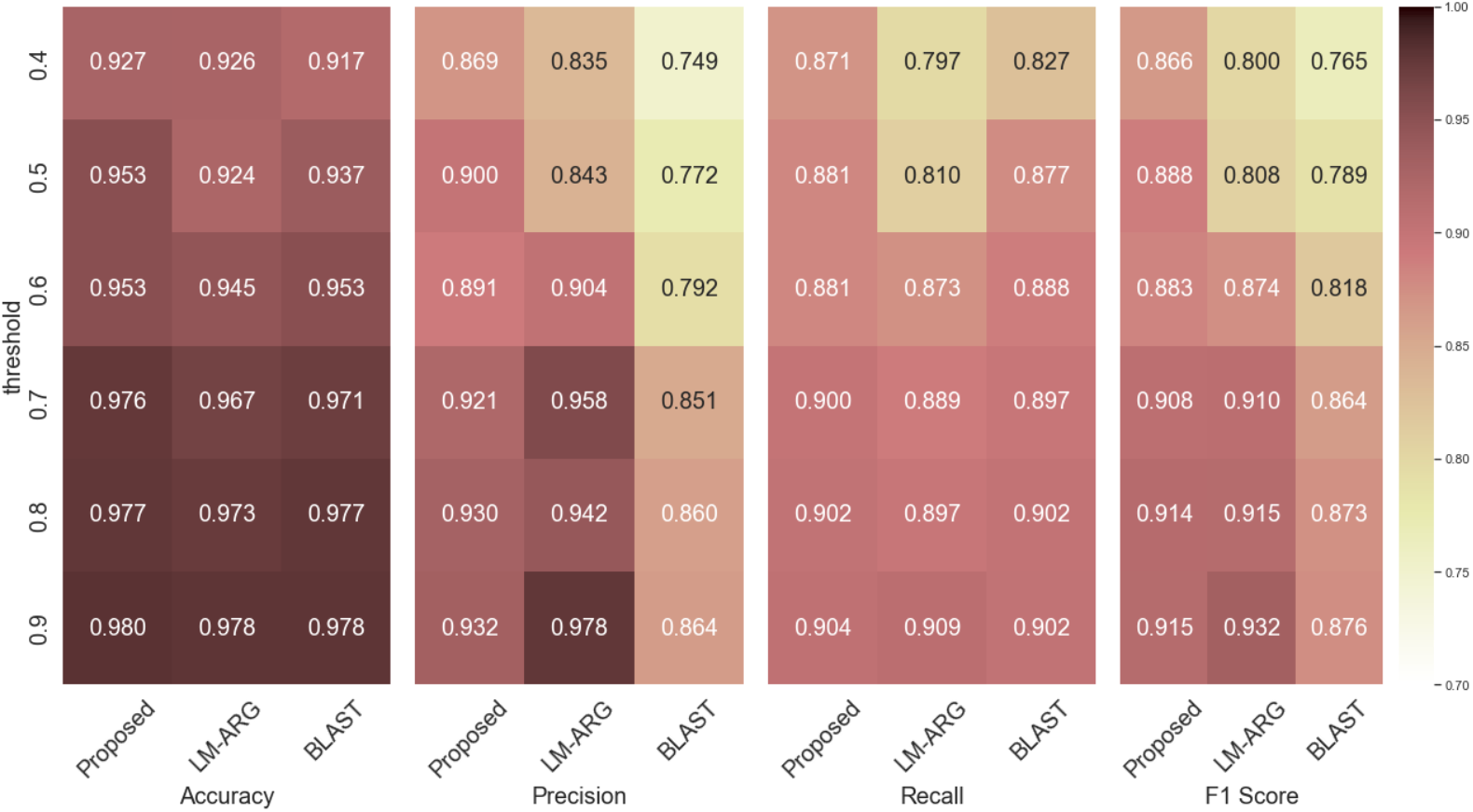
Performance comparison of the proposed method, LM-ARG, and BLASTn low homology datasets generated with varying sequence identity thresholds (see Section 2.1.2 for dataset generation details). with respcet to Accuracy, Precision, Recall, and F1 Score (see Section 2.5 for metric definitions). The y-axis indicates the sequence identity threshold used to create each low homology dataset, ranging from 0.4 to 0.9 in increments of 0.1.

### 3.2 Attention analysis

We selected four ARGs for attention analysis based on specific criteria. For macB and tetW, we first identified the two most prevalent families within each resistance mechanism. We then chose sequences that had at least one significant domain and contained a high number of domains (five or more). In the case of rpoB, our initial intention was to analyze this sequence alongside information on point mutations, so we selected a sequence with available point mutation data. Lastly, blaOXA-114s was chosen because beta-lactamases are highly prevalent and play a crucial role in the antibiotic inactivation mechanism. Supplementary Table S1 provides detailed information on the selected sequences, including the ARG family, HMD-ARG DB accession, resistance mechanism, and description of the ARGs.

#### 3.2.1 Relationship between attention and amino acid conservation

Previous studies have shown that in protein language models (pLMs) predicting solubility, highly conserved regions contribute to the prediction (Thumuluri *et al*., 2022). Motivated by this finding, we compared the attention values and conservation of amino acids for each residue in tetW, and found a significant correlation as in Supplementary Figure S5 (Spearman correlation coefficient 0.29, p=2.73E-14). Furthermore, amino acid residues were classified into three groups based on attention values: low, medium, and high, using the 33rd and 66th percentiles as thresholds (33rd percentile = -2.77e-4, 66th percentile = -8.72e-5). The Mann-Whitney U test (significance level p *<* 0.05/N, where N is the number of tests i.e., N = 3) was then used to compare the conservation score distributions between these groups. The results showed that the distribution of conservation scores for the ‘high’ group was significantly greater than the distributions for the ‘low’ and ‘medium’ groups (Medium vs High:p = 8.27e-14, Low vs High:p = 4.20e-12). However, no significant differences were found between the ‘low’ and ‘medium’ groups (Figure 3, p = 0.82). These findings suggest that attention in the proposed model recognizes highly conserved amino acid residues, but it cannot be concluded that amino acid residues with low conservation scores necessarily have low attention values.

**Fig. 3:**
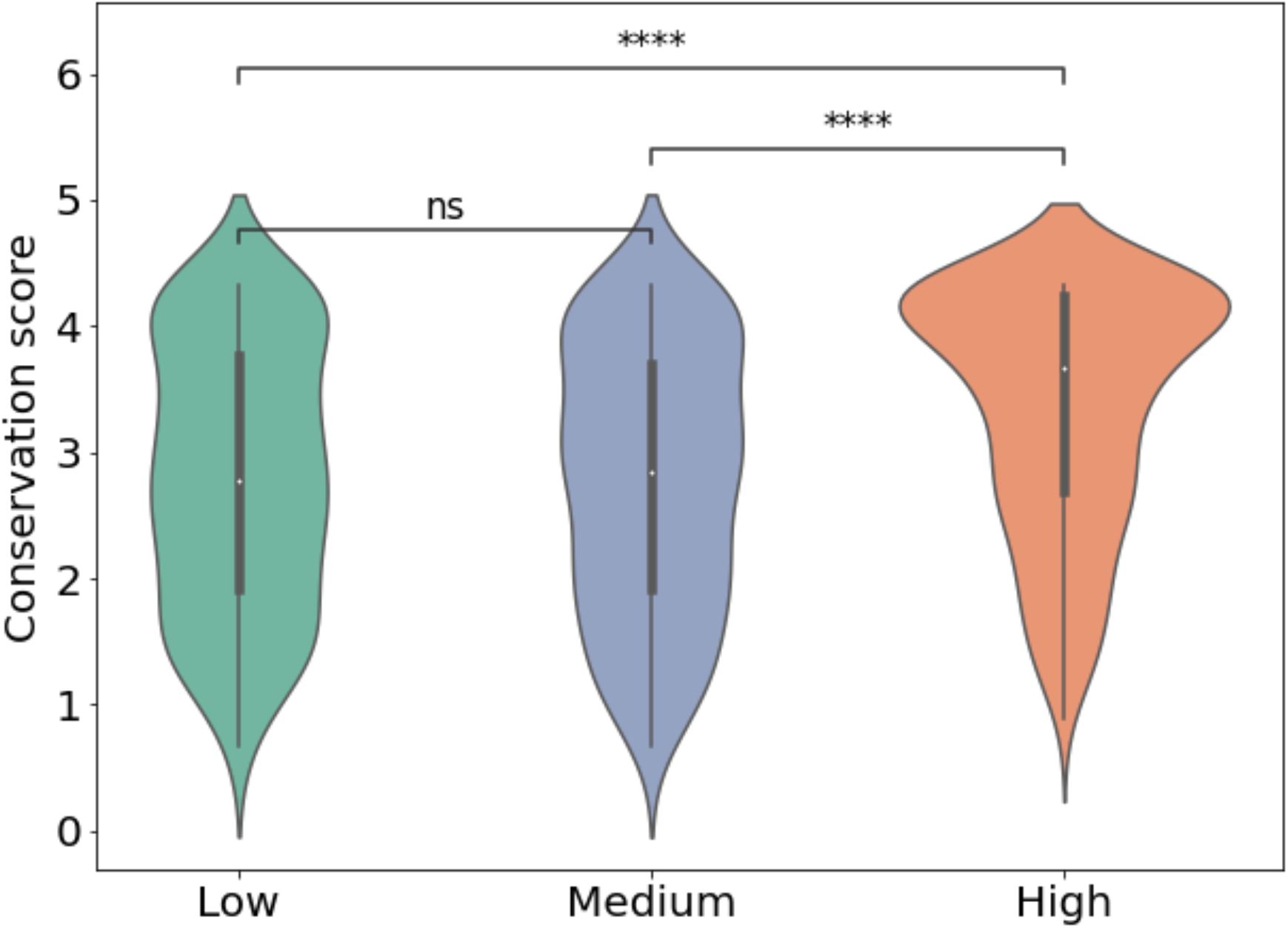
Relationship between attention values and amino acid conservation scores in tetW. Attention values were divided into three groups: Low (attention < 33rd percentile), Medium (33rd percentile < attention < 66th percentile), and High (attention > 66th percentile). The distribution of conservation scores was compared among these groups using the Mann-Whitney U test with Bonferroni correction (significance level: p < 0.05/3). “ns” indicates no significant difference (0.05/3 < p < 1), while “****” indicates a highly significant difference (p < 1.00e-04) between the conservation score distributions.

#### 3.2.2 Biological functions of attention-intensive sequence regions

Previous studies have shown that in BERT models pre-trained on antibody sequences, attention is concentrated on amino acid residues located in antigen-binding sites (Leem *et al*., 2022). We hypothesized that the attention in our proposed model would exhibit similar characteristics. In the following, we focus on four ARGs listed in Supplementary Table S1 for attention analysis, based on attention-intensive regions (cf. Section 2.6.2).

Table 2 (top) shows four attention-intensive regions in rpoB. Notably, domain PF04565, which represents RNA polymerase beta subunit domain 3, is known as the fork domain and is involved in binding to the target antibiotic rifampin (Alifano *et al*., 2015). On the other hands, Table 2 (bottom) presents the attention-intensive regions in tetW, with significantly high attention values observed in the GTP-binding domain. GTP-binding has been shown to play a crucial role in tetW-mediated antibiotic resistance (Chopra and Roberts, 2001); This finding is supported by the observation that mutations in the GTP-binding site altered the affinity for the target antibiotic (Chopra and Roberts, 2001). These results demonstrate that attention recognizes the antibiotic binding site in rpoB and the site that contributes to antibiotic resistance in tetW.

**Table 2:** Attention-intensive regions identified by InterProScan for rpoB and tetW. The table lists the significantly enriched InterPro signatures (p-value < 0.05 after Bonferroni correction) in the attention-intensive regions of each ARG. The columns show the anti-microbial registance (AMR) family, the accession number of the InterPro signature, the database from which the signature was derived, a brief description of the signature, and the p-value indicating the significance of the attention enrichment in the corresponding region.

| AMR Family | Accession | Database | Signature description | p-value |
| --- | --- | --- | --- | --- |
| rpoB | G3DSA:3.90.1100.10 | Gene3D | - | 2.49e-11 |
|  | G3DSA:3.90.1800.10 | Gene3D | RNA polymerase alpha subunit dimerisation domain | 4.86e-10 |
|  | G3DSA:3.90.1800.10:FF:000001 | FunFam | DNA-directed RNA polymerase subunit beta | 4.86e-10 |
|  | PF04565 | Pfam | RNA polymerase Rpb2, domain 3 | 1.94e-14 |
| tetW | PR00315 | PRINTS | GTP-binding elongation factor signature | 1.33e-08 |
|  | PTHR43261 | PANTHER | TRANSLATION ELONGATION FACTOR G-RELATED | 4.37e-05 |
|  | TIGR00231 | NCBIfam | JCVI: small GTP-binding protein domain | 1.12e-04 |

Next, we conducted more detailed attention analyses (Figure 4) ro rpoB and tetW. Figure 4a and 4c show the corresponding positions (top) and attention values (bottom) of the rpoB and tetW domains, respectively. These figures confirm that the regions where significant differences in the distribution of attention were identified show specifically high attention values. In addition, Figure4b and 4d illustrate the attention values mapped onto the predicted three-dimensional structures of rpoB and tetW using AlphaFold (Jumper *et al*., 2021; Varadi *et al*., 2022). Amino acids with high attention values, highlighted in red, are clustered around the purple binding molecules (rifampin in Figure4b, GTP in Figure4d) in three-dimensional spaces. All the figures also suggest that attention is concentrated on the rifampin or GTP binding site.

**Fig. 4:**
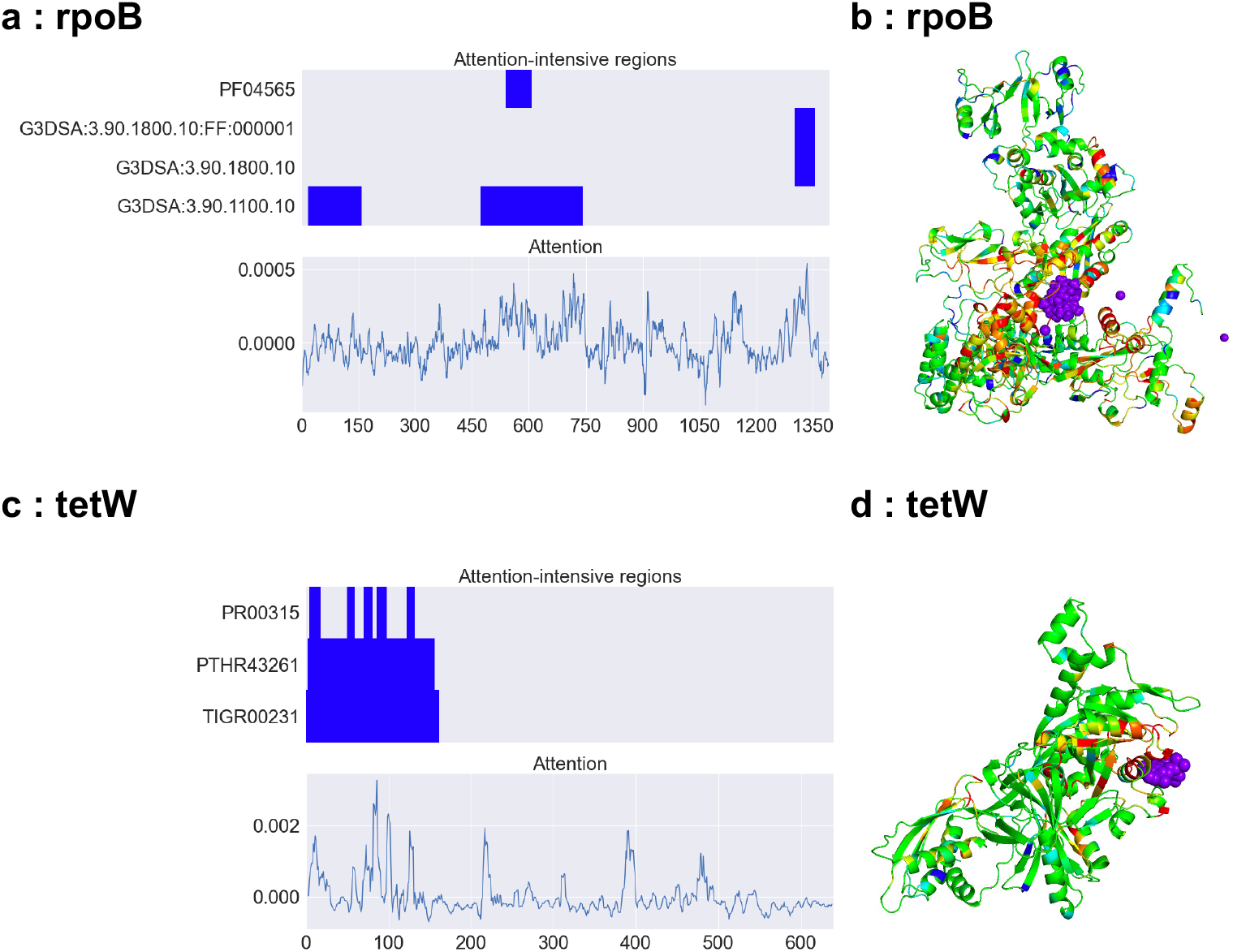
Attention analysis of rpoB and tetW. (a, c) Top: Attention-intensive regions (cf. Secton 3.2.2) for InterPro-identified regions in rpoB and tetW, respectively. Bottom: Smoothed attention values obtained by taking the average of five residues, with the target residue in the middle. The two graphs share an x-axis indicating the amino acid position. (b, d) Attention values mapped onto the 3D structures of rpoB (AlphaFold DB ID: AF-B4RQW2-F1) and tetW (AlphaFold DB ID: AF-A3R795-F1), respectively. Residues with high attention are colored red, while those with low attention are colored blue. The purple crystal structures represent rifampin and GTP, which were obtained from the crystal structures of homologous sequences (PDB IDs: 4kmu and 2hcj, respectively).

Supplementary Figure S6 and Table S2 indicate the results for BlaOXA-114a, suggesting that the class-D active site that hydrolyzes the target beta-lactam has higher attention values (Bauernfeind, 1986). Supplementary Figure S7 and Table S2 indicate the results for macB, an ABC transporter commonly found in antibiotic efflux, suggesting that significant attention concentrations are observed at the conserved site of the ABC transporter, the ATP-binding site. These results suggest that attention in the proposed method focuses on conserved regions that contribute to resistance.

Finally, we investigated whether the attention characteristics observed in individual ARGs are common across other sequences in the HMD-ARG DB. We identified sequence regions where attention was significantly concentrated and performed Gene Ontology (GO) enrichment analysis on these attention-intensive regions using Fisher’s exact test with Bonferroni correction for multiple testing. The results, as shown in Figure 5, revealed that GO terms such as ‘GTP binding’ and ‘GTPase activity’ were enriched in antibiotic target protection, while ‘ATP binding’ and ‘ATP hydrolysis activity’ were enriched in antibiotic target alteration. These findings suggest that the attention patterns observed in tetW and macB are consistent across other sequences. Furthermore, the enrichment of ‘antibiotic catabolic process’ in ‘antibiotic inactivation’ indicates that, similar to blaOXA-114s and rpoB, attention can identify sequence regions that bind and interact with the target antibiotic. Collectively, these results demonstrate that attention in our model recognizes interaction sites and highly conserved regions associated with target antibiotics across a wide range of ARGs in the HMD-ARG DB.

**Fig. 5:**
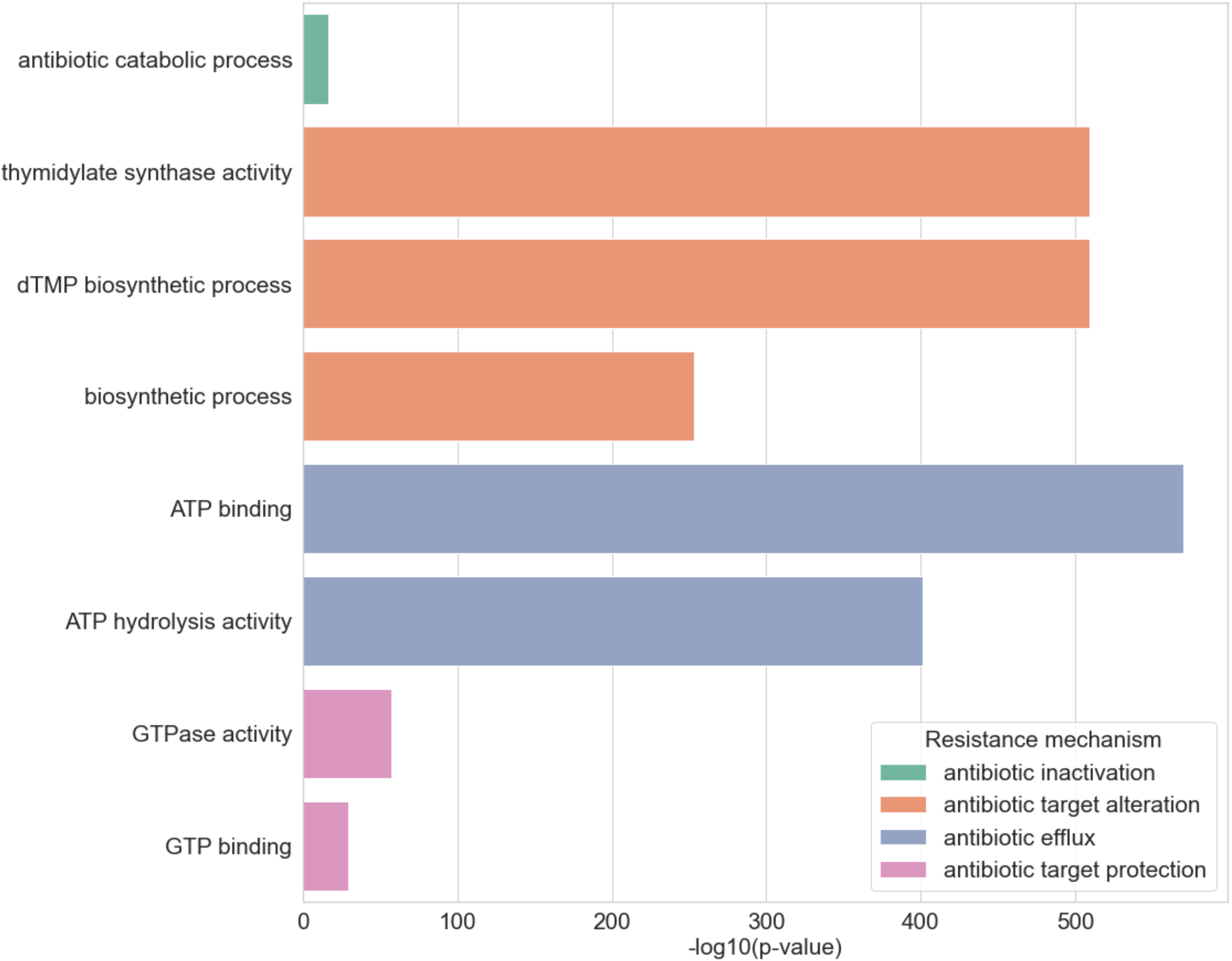
Gene Ontology terms enriched in attention-intensive regions within ARG sequences belonging to each resistance mechanism. GO terms annotated in over 150 attention-intensive regions were selected for visualization from all significantly enriched Gene Ontology terms using Fisher’s exact test. The x-axis represents the -log10(p-value) for each GO term, while the y-axis lists the GO terms. P-values were corrected for multiple testing using the Bonferroni method. The color of the bars indicates the type of resistance mechanism.

## 4 Discussion

In this study, we developed a novel computational approach to predict resistance mechanisms from ARG sequences using ProteinBERT, a pre-trained protein language model. Our proposed method outperformed existing methods across multiple datasets, including those with low sequence similarity to the training data (Section 3.1). This suggests that our model can accurately predict the resistance mechanisms of unknown ARGs, which is crucial for understanding the ARG reservoir and developing strategies to combat antibiotic resistance. Moreover, the attention analysis revealed that our model is interpretable and considers biologically relevant features, such as amino acid conservation and target binding sites when making predictions (Section 3.2). The attention analysis also suggests that our model may have the potential to recognize tertiary structures and point mutations, but further investigation is required to confirm these hypotheses. This interpretability is a significant advantage over existing methods and can provide valuable insights into the molecular basis of antibiotic resistance.

It is important to acknowledge the limitations of the current study, particularly in terms of data collection and analysis methods. Firstly, the reliability of the annotation methods used for training and evaluating datasets is questionable due to the lack of a unified definition and the scarcity of databases containing information on the nature of ARGs, such as resistance mechanisms. The inconsistency in annotation methods, with some based on the AMR Family and others relying on the predicted results of existing methods, further compounds this issue. The credibility of these methods remains uncertain, and the development of a more accurate and uniform database would greatly assist in predicting and analyzing resistance mechanisms. Secondly, our analysis may have overlooked unknown attention-intensive regions, as we focused on identifying regions of attention concentration from known databases. This approach fails to identify attention-intensive regions that are not present in the database and may miss unknown functional sites related to resistance mechanisms. Improved methods for analyzing attention could potentially reveal important sequence regions that have not yet been discovered, providing deeper insights into the molecular basis of antibiotic resistance.

Despite these limitations, our study demonstrates the potential of using pre-trained protein language models for predicting and understanding ARG resistance mechanisms, with several promising applications. The attention analysis results, which identify conserved regions and binding sites associated with antibiotic resistance, could guide researchers in designing novel antibiotics that target these specific areas, potentially leading to the development of more effective drugs that can overcome existing resistance mechanisms. Furthermore, by leveraging the model’s ability to accurately predict resistance mechanisms, even for ARGs with low sequence similarity to known sequences, it may be possible to develop rapid diagnostic tools for the early detection and monitoring of antibiotic-resistant bacteria. These tools could help healthcare professionals identify resistant strains more quickly, allowing for the timely implementation of appropriate treatment strategies and infection control measures, ultimately contributing to the global effort in combating antibiotic resistance.

## Acknowledgments

We would like to express our sincere appreciation to the members of the Hamada Laboratory, especially Drs. Hitoshi Iuchi and Junna Kawasaki. We extend our gratitude to the participants of the 6th Tokyo Bioinformatics Meeting for their invaluable insights.

