## Supplementary material for "Prediction of antibiotic resistance mechanisms using a protein language model": Supplymentaly File

### Supplementary materials for
### “Prediction of antibiotic resistance mechanisms using a protein language model”

Kanami Yagimoto, Shion Hosoda, Miwa Sato and Michiaki Hamada

### 1 Supplementary Figures and Tables

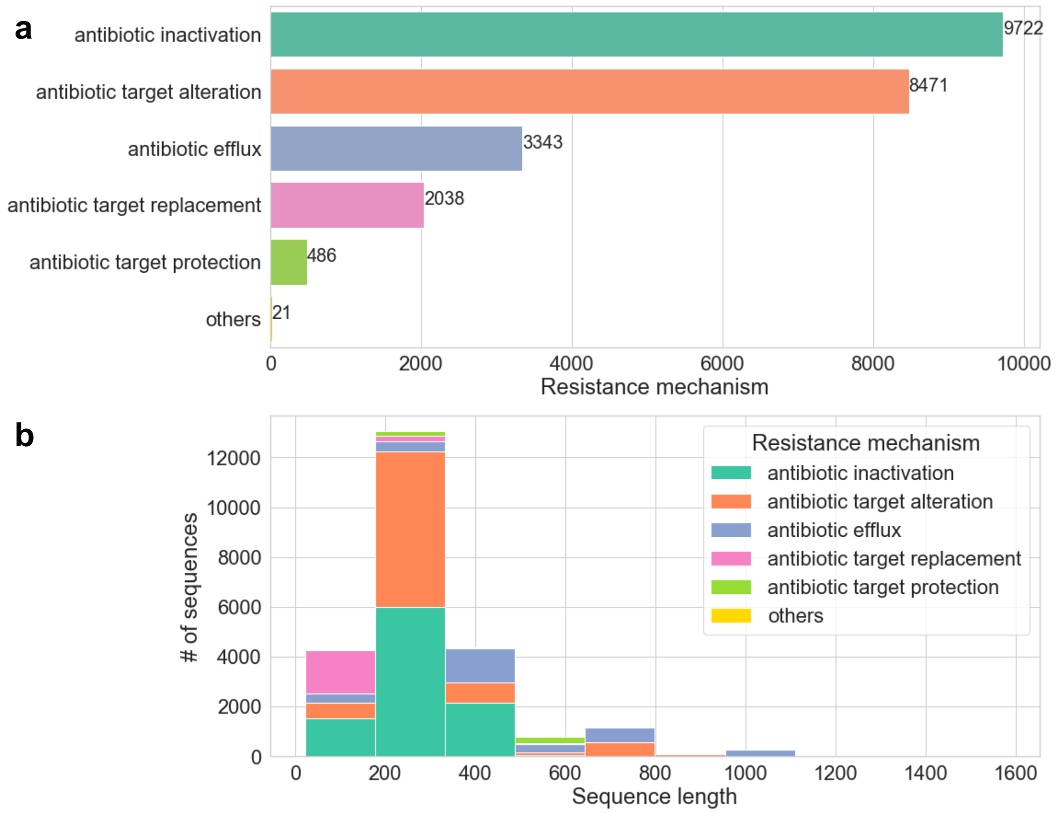


Fig. S1: Overview of the HMD-ARG DB and Low Homology Dataset. Both datasets contain the same ARGs but differ in sequence similarity between the training and test sets. a) Distribution of ARGs across different resistance mechanisms. The bar plot shows the number of ARGs associated with each resistance mechanism. b) Distribution of ARG sequence lengths. The histogram illustrates the frequency of ARGs with respect to their sequence length in amino acids.

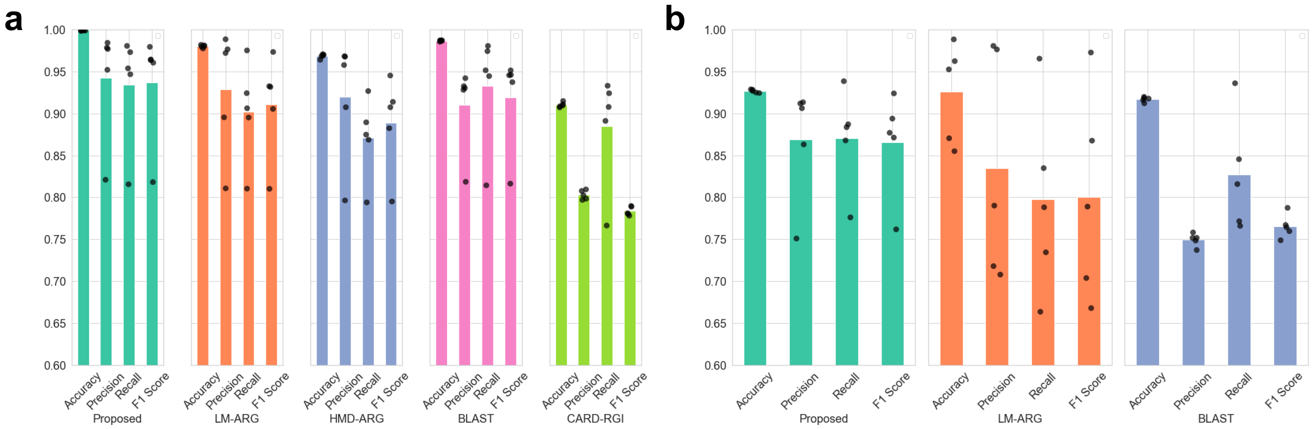


Fig. S2: Performance metrics for each resistance mechanism and their macro average, obtained from 5-fold cross-validation on a) HMD-ARG DB and b) Low Homology Dataset. The black dots represent the values for each iteration of the 5-fold cross-validation, while the bar plots show the average values across all iterations. The metrics evaluated include Accuracy, Precision, Recall, and F1-score. The macro average provides an overall assessment of the model's performance across all resistance mechanisms.

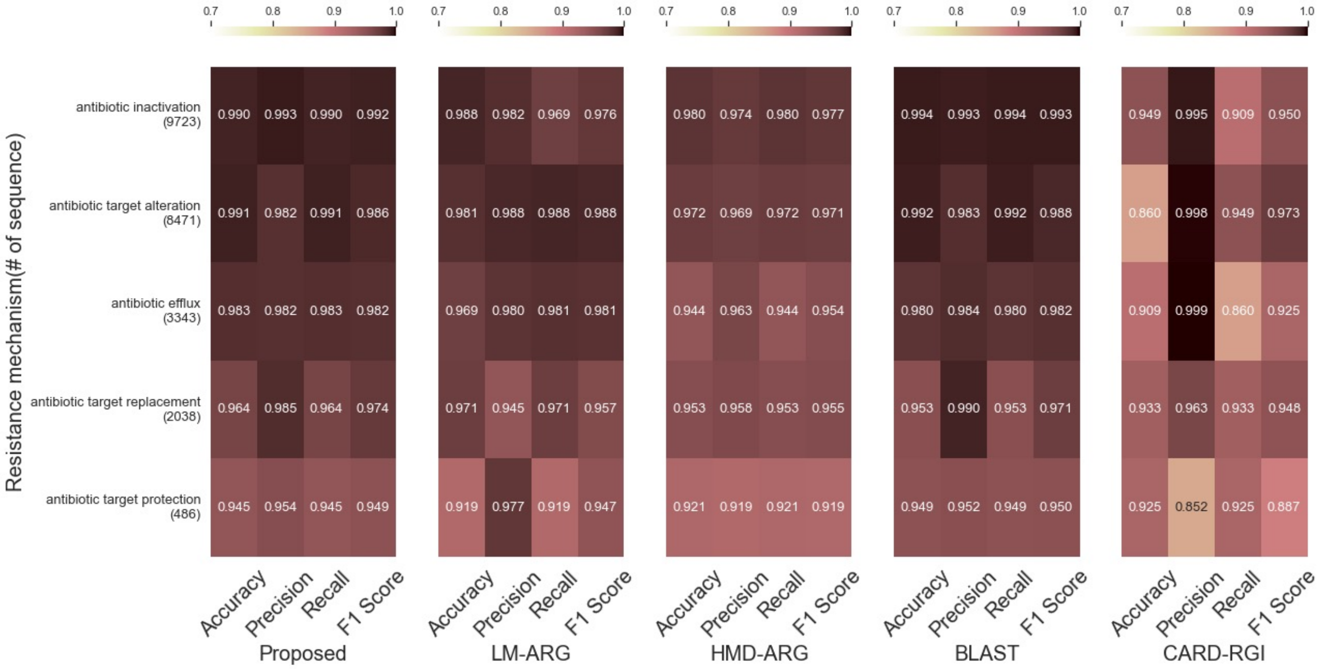


Fig. S3: Performance comparison of the proposed method with existing methods (LM-ARG, HMD-ARG, BLAST, and CARD-RGI) for each resistance mechanism in the HMD-ARG DB. The number of sequences associated with each resistance mechanism is indicated in parentheses on the y-axis. The bar plots represent the F1-score, a balanced measure of Precision and Recall, for each method and resistance mechanism.

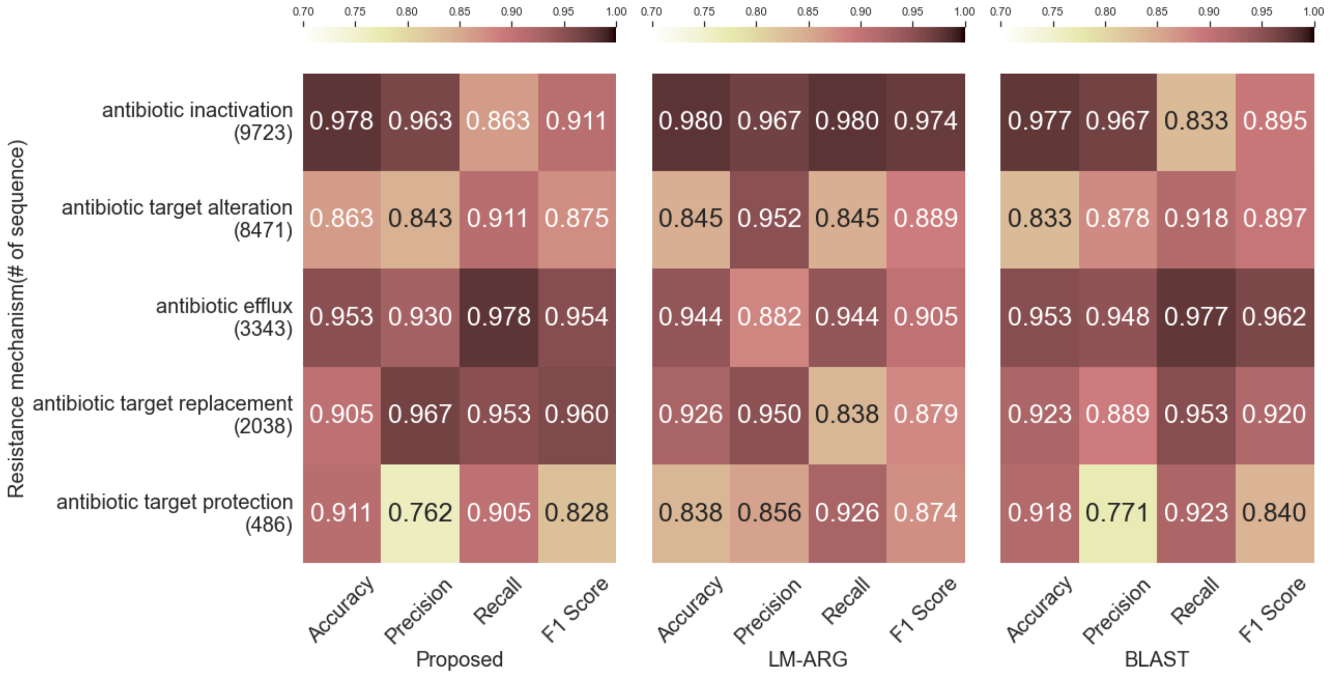


Fig. S4: Performance comparison of the proposed method with LM-ARG and BLAST for each resistance mechanism in the Low Homology Dataset (with 0.4 of the similarity threshold). The number of sequences associated with each resistance mechanism is indicated in parentheses on the y-axis. The bar plots represent the F1-score, a balanced measure of Precision and Recall, for each method and resistance mechanism.

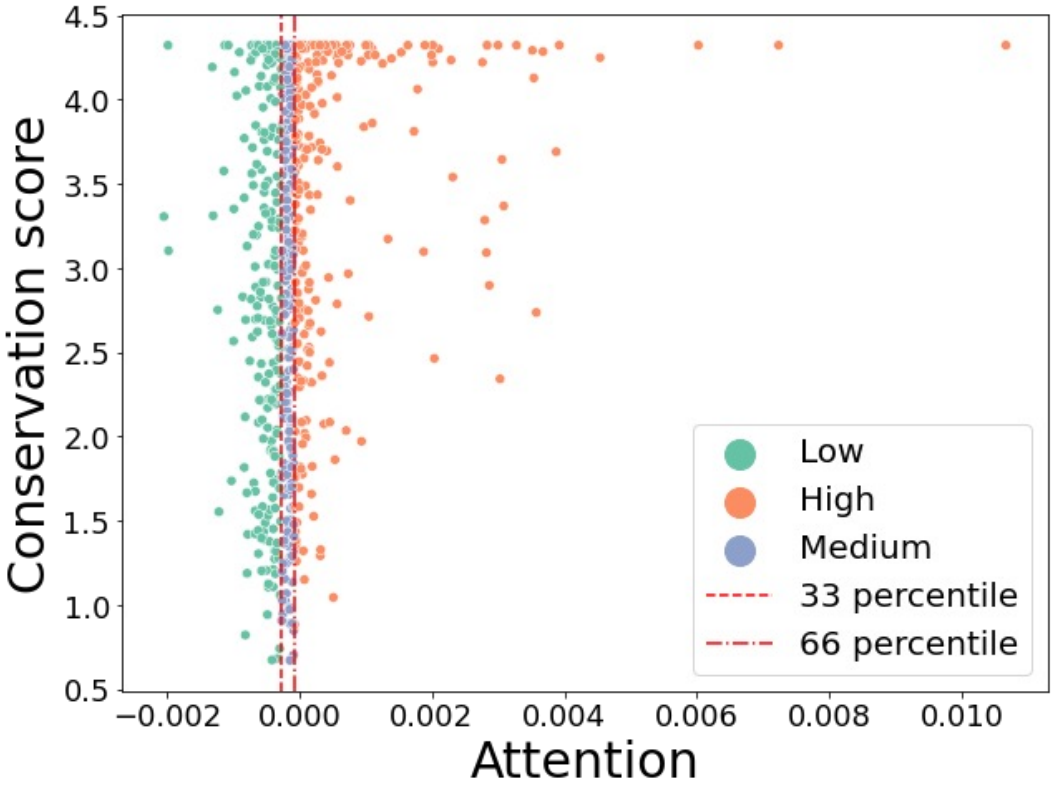


Fig. S5: The scatterplot of attention (x-axis) and conservation score (y-axis) in tetW. Amino acid residues of tetW were divided into three groups according to their attention values, minimum 33 percentile,33 percentile 66 percentile, and 66 percentile. The red vertical lines in this figure represented the 33 and 66 percentiles. The groups were then named Low/Medium/High and each plot in this figure was colored according to their groups.

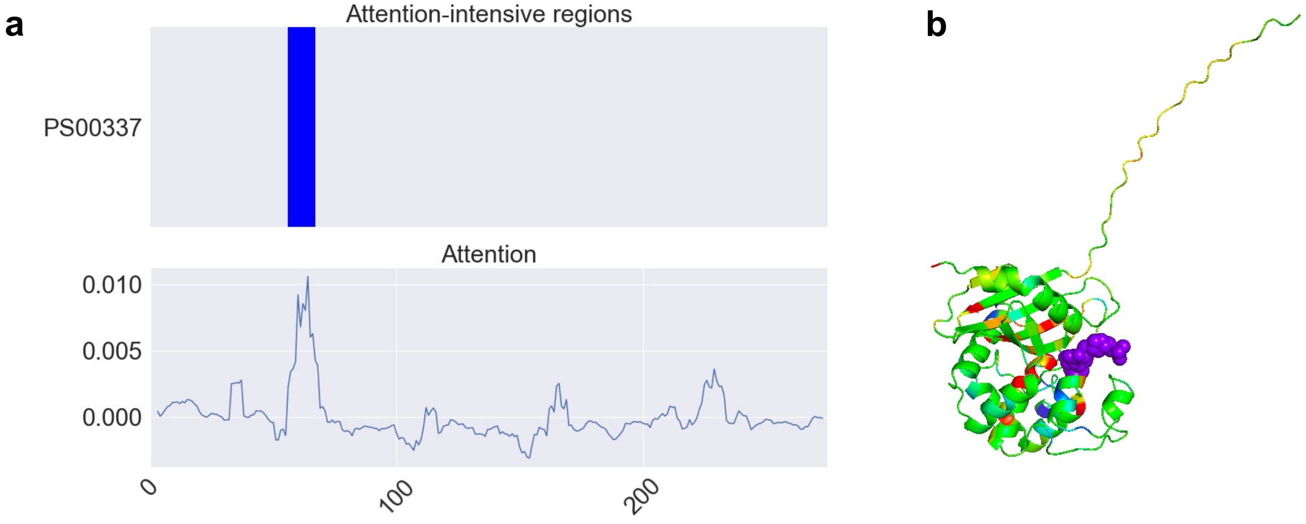


Fig. S6: Attention analysis of the blaOXA-114a gene. a) Top: Attention-intensive regions in blaOXA-114a. Bottom: Smoothed attention values obtained by calculating the average attention of each amino acid residue and its two neighboring residues on either side. The x-axis indicates the amino acid position in blaOXA-114a, shared by both plots. b) Attention values mapped onto the predicted 3D structure of blaOXA-114a (AlphaFold DB ID: AF-U3N8W9-F1). The purple crystal structure represents the target antibiotic, which was obtained from the crystal structure of a homologous sequence (PDB ID: 3isg).

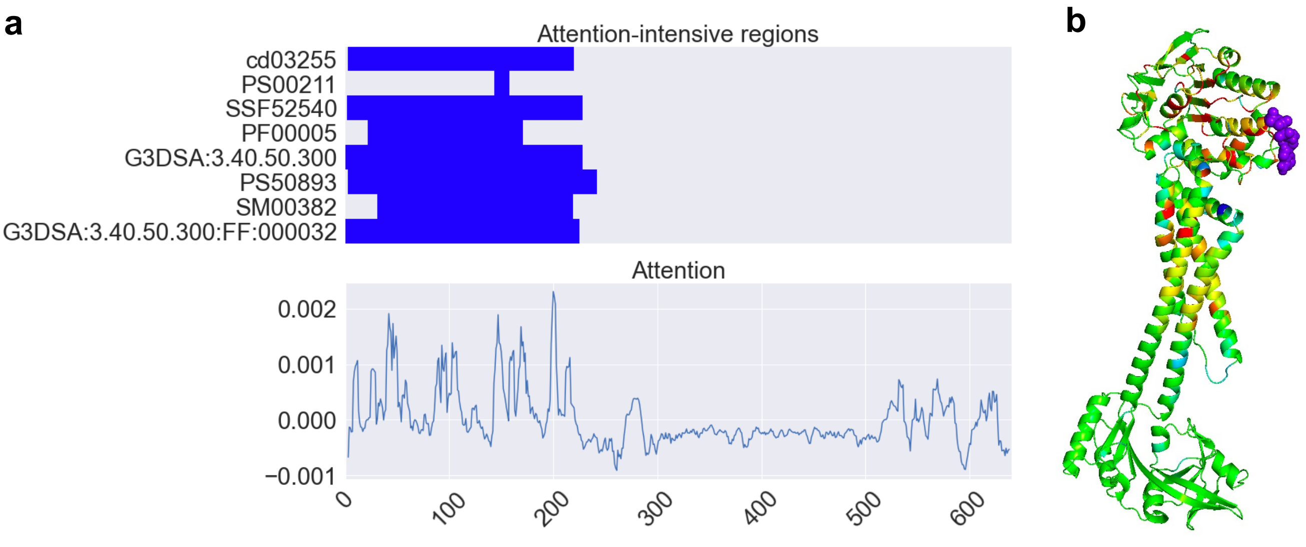


Fig. S7: Attention analysis of the macB gene. a) Top: Attention-intensive regions in macB. Bottom: Smoothed attention values obtained by calculating the average attention of each amino acid residue and its two neighboring residues on either side. The x-axis, shared by both plots, indicates the amino acid position in macB. b) Attention values mapped onto the predicted 3D structure of macB (AlphaFold DB ID: AF-A0A011P660-F1). The purple crystal structure represents ATP, obtained from the crystal structure of a macB homolog sequence (PDB ID: 5lil).

Table S1: A list of target sequences of attention analysis.

| subclass | HMD-ARG DB Accession | Resistance mechanism | Description |
|---|---|---|---|
| rpoB | NP_273190.1 | antibiotic target alteration | rifamycin-resistant beta-subunit of RNA polymerase (rpoB) |
| tetW | ABN80187 | antibiotic target protection | tetracycline-resistant ribosomal protection protein |
| blaOXA-114s | U3N8W9 | antibiotic inactivation | OXA beta-lactamase |
| macB | A0A011P660 | antibiotic efflux | ATP-binding cassette (ABC) antibiotic efflux pump |

Table S2: Interpro-registered regions in blaOXA-114a and macB with significantly higher attention values compared to other regions. The table includes the AMR family, accession number, database, signature description, and p-value for each significant region.

| AMR Family | Accession | Database | Signature description | p-value |
|---|---|---|---|---|
| bla-OXA-114s | PS00337 | ProSitePatterns | Beta-lactamase class-D active site. | 1.13e-03 |
| macB | G3DSA:3.40.50.300 | Gene3D | - | 3.62e-10 |
| | G3DSA:3.40.50.300:FF:000032 | FunFam | Export ABC transporter ATP-binding protein | 1.28e-10 |
| | PF00005 | Pfam | ABC transporter | 1.22e-08 |
| | PS00211 | ProSitePatterns | ABC transporter family signature. | 5.22e-07 |
| | PS50893 | ProSiteProfiles | ATP-binding cassette, ABC transporter-type domain profile. | 1.11e-08 |
| | SM00382 | SMART | AAA_5 | 2.23e-11 |
| | SSF52540 | SUPERFAMILY | P-loop containing nucleoside triphosphate hydrolases | 8.15e-11 |
| | cd03255 | CDD | ABC_MJ0796_LolCDE_FtsE | 1.79e-11 |

### 2 Supplementary Text

#### 2.1 HMD-ARG DB

First, the genome-encoded datasets were translated into protein sequences. The resulting protein sequences were then merged, and duplicates were removed using BLASTP and CD-HIT with a sequence identity threshold of 100%. This process yielded a total of 24,082 unique ARGs. Next, resistance mechanism labels were assigned to each ARG using the CARD ontology. The labels included antibiotic efflux, antibiotic inactivation, antibiotic target alteration, antibiotic target protection, antibiotic target replacement, and others. The assignment was performed using the best hit strategy with BLASTP and a cut-off E-value of 1e-20. For sequences that could not be assigned a label using this approach, manual curation was performed.

#### 2.2 Fine-tuning for pre-trained model

The fine-tuning of the proposed method was carried out in three stages to optimize its performance:

1. First epochs: In this stage, all layers of the pre-trained model were frozen, and only the newly added fully connected layers were trained for up to 40 epochs. The learning rate was optimized epoch by epoch, starting from an initial learning rate of 1e-02.

2. Next epochs: Following the first stage, all layers were unfrozen, and the entire model was trained for an additional 40 epochs. The learning rate was optimized epoch by epoch, starting from an initial learning rate of 1e-04.

3. Final epoch: In the last stage, the model was trained for a final epoch using sequences longer than the maximum sequence length (1,024 amino acids) and a learning rate of 1e-05. Throughout all epochs, the learning rate was adjusted using the ReduceLROnPlateau callback from Keras, with the following parameters: patience=1, factor=0.25, and min_lr=1e-05. This callback reduces the learning rate when the validation loss plateaus, allowing for more refined adjustments to the model's weights.

Additionally, early stopping was implemented using the EarlyStopping callback from Keras, with the following parameters: patience=2 and restore_best_weights=True. This callback monitors the validation loss and stops the training process if no improvement is observed for a specified number of epochs, preventing overfitting and ensuring that the best-performing model weights are retained.

It is important to note that the GO annotations of the training sequences were not utilized during the fine-tuning process, as the focus was on predicting antibiotic resistance mechanisms solely based on the protein sequences.

To evaluate the model's performance and generalization ability, a validation dataset was created by randomly selecting 10% of the training data. This validation set was used to monitor the model's performance during training and to guide the application of early stopping.

#### 2.3 Conservation Score Calculation for Attention Analysis

To calculate the conservation score for each amino acid position in the attention analysis, the following steps were performed (Figure S8):

1. Homologous sequence retrieval: PSI-BLAST (Altschul *et al.*, 1997; Schäffer *et al.*, 2001) was employed to search for homologous sequences in the RefSeq database (O'Leary *et al.*, 2016), a comprehensive, non-redundant protein sequence database. The search was conducted for 5 iterations with an E-value threshold of 0.0001, and the top 500 homologous sequences were retrieved for each input sequence.

2. Multiple sequence alignment (MSA): The obtained homologous sequences were aligned using ClustalW (Thompson *et al.*, 1994; Larkin *et al.*, 2007) to generate a multiple sequence alignment (MSA). This alignment enables the identification of conserved regions and residues across the homologous sequences.

3. Conservation score calculation: For each position in the MSA, the frequency of occurrence of each amino acid was determined. The conservation score was then calculated as the Kullback-Leibler (KL) divergence between the observed amino acid distribution at each position and a uniform distribution. The KL divergence quantifies the difference between two probability distributions, with higher values indicating a greater deviation from the uniform distribution and, thus, a higher level of conservation.

The conservation scores calculated using this method were then utilized in the attention analysis to examine the relationship between the attention values assigned by the proposed model and the evolutionary conservation of amino acid positions in the input sequences.

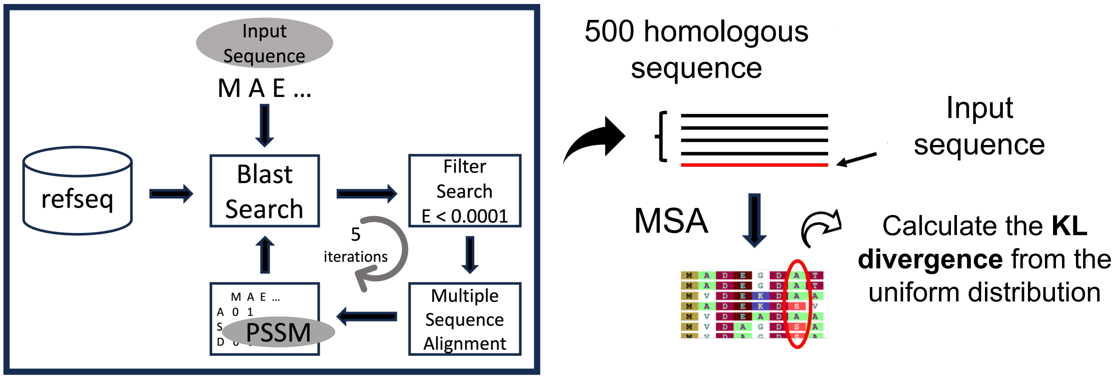


Fig. S8: Method for calculating the conservation score used in the attention analysis
